# Prophylactic low-dose, bi-weekly benznidazole treatment fails to prevent *Trypanosoma cruzi* infection in dogs under intense transmission pressure

**DOI:** 10.1101/2022.07.25.501361

**Authors:** Juan M. Bustamante, Angel M. Padilla, Brooke White, Lisa D. Auckland, Rachel E. Busselman, Stephanie Collins, Elizabeth L. Malcolm, Briana F. Wilson, Ashley B. Saunders, Sarah A. Hamer, Rick L. Tarleton

## Abstract

*Trypanosoma cruzi* naturally infects a wide variety of wild and domesticated mammals, in addition to humans. Depending on the infection dose and other factors, the acute infection can be life-threatening, and in all cases, the risk of chagasic heart disease is high in persistently infected hosts. Domestic, working, and semi-feral dogs in the Americas are at significant risk of *T. cruzi* and in certain settings in the southern United States, the risk of new infections can exceed 30% per year, even with the use of vector control protocols. In this study, we explored whether intermittent low-dose treatment with the trypanocidal compound benznidazole (BNZ) during the transmission season, could alter the number of new infections in dogs in an area of known, intense transmission pressure. Preliminary studies in mice suggested that twice-weekly administration of BNZ could prevent or truncate infections when parasites were delivered at the mid-point between BNZ doses. Pre-transmission season screening of 126 dogs identified 53 dogs (42.1%) as *T. cruzi* infection positive, based upon blood PCR and Luminex-based serology. Serial monitoring of the 67 uninfected dogs during the high transmission season (May to October) revealed 15 (22.4%) new infections, 6 in the untreated control group and 9 in the group receiving BNZ prophylaxis, indicating no impact of this prophylaxis regimen on the incidence of new infections. Although these studies suggest that rigorously timed and more potent dosing regimen may be needed to achieve an immediate benefit of prophylaxis, additional studies would be needed to determine if drug prophylaxis reduced disease severity despite this failure to prevent new infections.

**Author Summary:** *Trypanosoma cruzi*, the parasite that causes Chagas disease, circulates extensively in the southern U.S. and working dog populations in south-central Texas are at very high risk of infection, and morbidity and early mortality due to this infection. In this study, we used low level administration of an FDA-approved drug during the transmission season to attempt to prevent new infections in these dogs. Although that effort failed, the study revealed new information about the transmission dynamics and generation of antibody responses and immune control of infection in this high-transmission setting.

## Introduction

Chagas disease, caused by the protozoan parasite *Trypanosoma cruzi*, is a problem for human and animal health across the Americas where triatomine vectors are endemic. *T. cruzi* is predominantly transmitted in the feces of infected triatomines through contact with wounds or mucous membranes or ingestion of infected insects or fecal material [1]. Oral transmission is thought to be the most important route in domestic dogs and is a highly efficient mode of transmission [2-5].

Dogs are important to study in the context of Chagas disease for at least three reasons: i) they experience similar disease progression to humans, so are useful to study as a model for human disease especially in considering treatments; ii) they share exposure to vectors with humans in domestic and peridomestic environments and therefore may serve as sentinels for human health risk [6-8]; and iii) canine Chagas disease is an increasingly recognized problem in veterinary medicine, especially in the southern U.S. [9], leading to dog mortality [10].

Large multi-dog kennels in central and south Texas have emerged as settings with particularly high transmission risk, with a recent study documenting incidence of over 30 infections per 100 dogs per year [11], despite variable vector control efforts in and around the kennels. Although some therapeutic regimens have shown promise in reducing disease impact in dogs infected with *T. cruzi* [12], no therapeutics have been evaluated as potential preventatives of *T. cruzi* infection in dogs.

Here we trialed a novel approach for preventing new infections in dogs within multi-dog kennel environments with a history of triatomine occurrence and canine Chagas disease. Disappointingly, prophylaxis during the period of peak adult vector activity [13] with benznidazole (BNZ), an FDA-approved drug used in the treatment of human *T. cruzi* infection, had no impact on the incidence of new infections in this setting, However the long-term impact of prophylaxis on clinical disease was not monitored. Further, this study design provides a model for future evaluation of different prophylactic regimens (higher dose; treatments earlier in the season) in high transmission settings.

## Material and Methods

### Study Design

For the studies in mice, female mice, 8-12 weeks old were used for infections throughout the study. Sample sizes were determined based upon our knowledge of heterogeneity in parasite burden during *T. cruzi* infection and published reports using similar experimental strategies. Data collected were included if productive *T. cruzi* infection was established (visualized by luciferase imaging or flow cytometry by detection of *T. cruzi*-tetramer specific CD8^+^ T cells). The investigators were not blinded during the collection or analysis of data and mice were randomly assigned to treatment groups prior to the start of each experiment.

For studies in dogs, we formed a small network of kennels in central and south Texas with a history of triatomine vector occurrence and canine Chagas disease with owners who were willing to participate in the study. Kennels are identified through (i) canine patients with Chagas disease presenting to the cardiology unit at the TAMU VMTH and (ii) the TAMU Kissing Bug Community Science program, in which many dog owners submit triatomines collected from large kennel environments. At these large kennels, dogs are primarily bred and trained to aid hunting parties. Approximately 40-80 dogs reside within each kennel, and the predominant breeds include Belgian Malinois, Brittany spaniels, English pointers, German shorthaired pointers, Labrador retrievers, and hound dogs. Dogs >3 months of age, including males and females, were eligible for enrollment. Dogs identified after blood screening as seronegative based on multiplex serology and PCR-negative were randomly assigned to treatment groups (Table S1). At each sampling event, up to 6ml of blood was collected from dogs via jugular venipuncture into clot activator tubes which were shipped on ice overnight to the laboratory after which serum was separated and used for antibody tests, and the clot was used for DNA extractions and PCR testing.

The dog owners/kennel managers gave prophylaxis by placing the encapsulated BNZ in the back of the dog’s mouth and encouraging swallowing; in some cases, the capsules were placed into cheese or other food for the dog to consume. Neither the investigators, private veterinarian providing care to the dogs in the field, nor the dog owners were blinded to the treatment groups and no placebo was given to control dogs.

### Mice, parasites and infections

C57BL/6J (Stock No:000664) mice (C57BL/6 wild-type) were purchased from The Jackson Laboratory (Bar Harbor, ME) and C57BL/6J-IFN-gamma knockout mice (also known as B6.129S7-Ifngtm1Ts/J; The Jackson Laboratory stock No 002287) were bred in-house at the University of Georgia Animal Facility. All the animals were maintained in the University of Georgia Animal Facility under specific pathogen-free conditions. *T. cruzi* tissue culture trypomastigotes of the Colombiana strain co-expressing firefly luciferase and tdTomato reporter proteins, generated as described previously [14, 15], were maintained through passage in Vero cells (American Type Culture Collection (Manassas, VA)) cultured in RPMI 1640 medium with 10% fetal bovine serum at 37°C in an atmosphere of 5% CO_2_. Mice were infected intraperitoneally with 10^3^ tissue culture trypomastigotes of *T. cruzi*.

### Drug treatment and in vivo imaging

Mice were treated with benznidazole twice weekly at a 100mg/kg/day and infected midway during the 3^rd^ week of treatment (Figure 1). Benznidazole (BNZ – Elea Phoenix, Buenos Aires, Argentina) was prepared by pulverization of tablets followed by suspension in an aqueous solution of 1% sodium carboxymethylcellulose with 0.1% Tween 80 and delivered orally by gavage at a concentration dosage of 100mg/kg of body weight. Each mouse received 0.2 ml of this suspension. Luciferase-expressing parasites were quantified in mice by bioluminescent detection. Mice were injected intraperitoneally with D-luciferin (150 mg/kg; PerkinElmer, Waltham, MA) and anesthetized using 2.5% (vol/vol) gaseous isofluorane in oxygen prior to imaging on an IVIS Lumina II imager (Xenogen, Alameda, CA), as previously described [16]. Quantification of bioluminescence and data analysis was performed using Aura Imaging Software version 4.0.7 (Spectral Instruments Imaging, Tucson, AZ).

**Figure 1.**
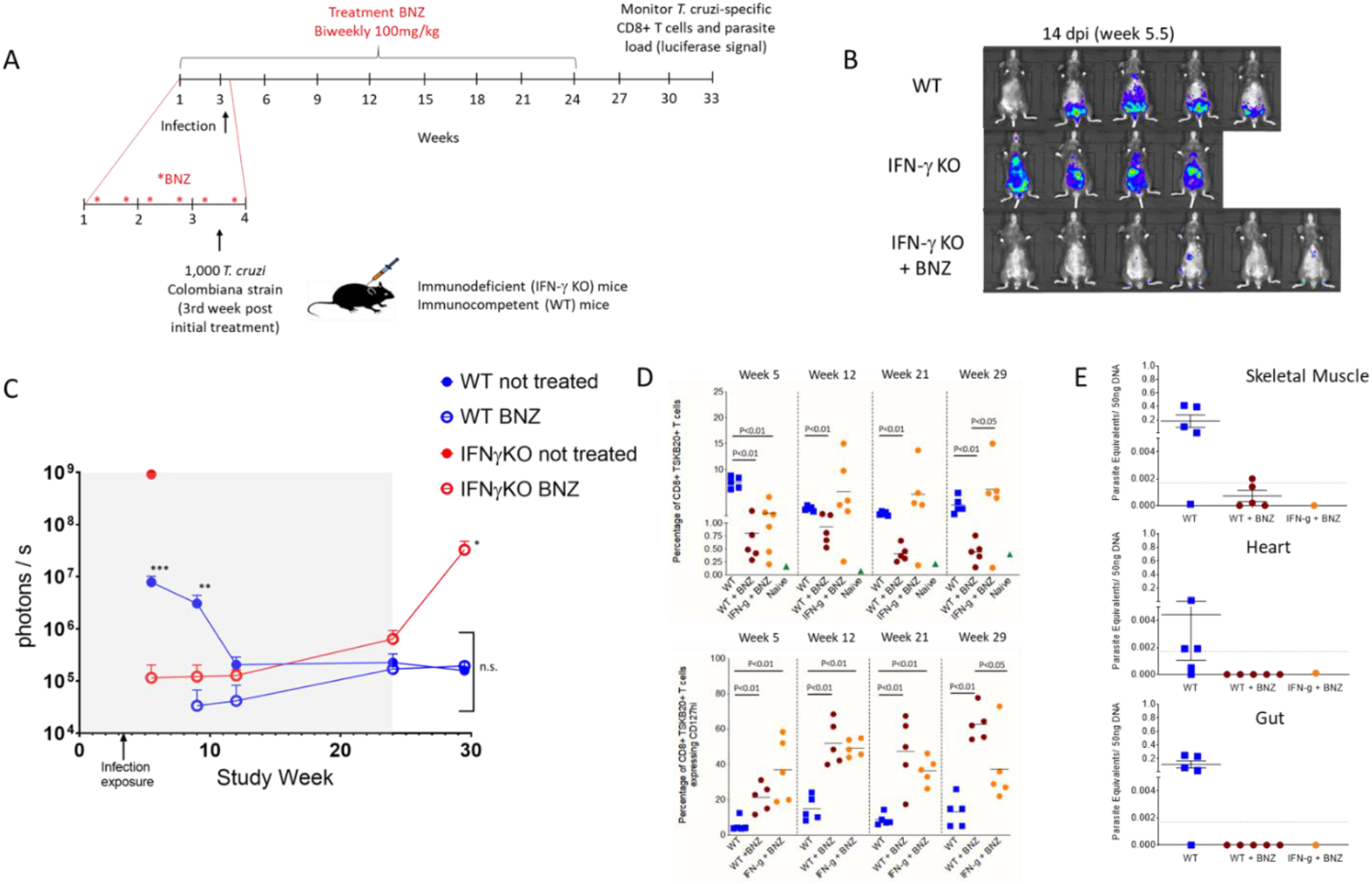
**(A)** Schematic representation of prophylactic treatment, infection, and monitoring of mice. WT and IFN*γ* deficient mice were treated biweekly with 100mg/kg of BNZ over 24 weeks. After the second week of treatment, mice were infected (5-6 animals/group) intraperitoneally with 10^3^ trypomastigotes of the Luciferase-expressing Colombiana strain of *T. cruzi*. Control groups of mice were infected and left untreated. **(B)** Representative images showing bioluminescent signal in mice (ventral view) at 14 days post-infection exposure. **(C)** Parasite bioluminescence quantification following D-luciferin injection was measured at the indicated times in the study period. Each data point represents the mean of bioluminescence from 3-6 mice expressed on a logarithmic scale as mean photons/second ±/-standard error of the mean. Grayed box marks the prophylaxis period. * p<0.03 relative to both WT groups; ** p<0.008 relative to both BNZ-treated groups; *** p<0.005 relative to BNZ-treated IFN*γ* KO group; n.s. bracketed data points are not statistically different. **(D)** Percentage of CD8^+^ blood lymphocytes specific for the *T. cruzi* TSKb20 epitope (top) and expressing the T cell central memory marker CD127 (bottom) during the study course **(E)** *T. cruzi* DNA determined by quantitative real time polymerase chain reaction in the skeletal muscle, heart, and intestine of untreated and BNZ-treated mice at week 33 of the study (∼30 weeks post-infection).

### T-cell phenotyping

Mouse peripheral blood was obtained, processed and analyzed as previously described [15, 17]. Whole blood was incubated with a major histocompatibility complex I (MHC I) tetramer containing the *T. cruzi* transialidase TSKB20 peptide (ANYKFTLV/Kb) labeled with BV421 (Tetramer Core Facility at Emory University, Atlanta, GA) and the following labeled antibodies: anti-CD8 FITC, anti-CD4 APC EF780, anti-CD127 PE (BD Bioscience, San Jose, CA). At least 500,000 cells were acquired using a CyAn ADP flow cytometer (Beckman Coulter, Hialeah, Florida) and analyzed with FlowJo software v10.6.1 (Treestar, Inc., Ashland, OR).

### Quantitative Polymerase Chain Reaction

For determination of tissue parasite load in mice, samples of skeletal muscle, heart and intestine were collected at necropsy and processed for quantification of *T. cruzi* DNA by real-time polymerase chain reaction (qPCR) as previously described [15, 17, 18]. The limit of detection was set at the lowest standard of 0.0017 parasite equivalents per 50 ng of DNA. For detection of *T. cruzi* infection in dogs, DNA was extracted from 250uL of the buffy coat fraction of EDTA-treated blood using the E.Z.N.A Tissue DNA kit (Omega Bio-Tek, Norcross, GA, USA) according to the manufacturer’s protocol except 50 μL of elution buffer was used. *T. cruzi-*negative controls (phosphate buffered saline) were included in the DNA extractions. Samples were tested using qPCR for the presence of *T. cruzi* satellite DNA using the Cruzi 1/2 primer set and Cruzi 3 probe in a real-time assay, which amplifies a 166-bp segment of a repetitive nuclear DNA [19, 20] as previously described [21] A sample was considered *T. cruzi-*positive if the Ct value was <35 [22].

### Multiplex serology

Luminex-based multiplex serological assays were performed as previously described [23, 24]. For a number of smaller proteins, fusions of up to 2 individual genes are employed in order to expand the array of antibodies being detected. (TritrypDb.org identifiers: Antigen **1** (Tc1) = fusion of TcBrA4_0116860 and TcYC6_0028190; **2** (Tc2) = fusion of TcBrA4_0088420 and TcBrA4_0101960; **3** (Tc3) = fusion of TcBrA4_0104680 and TcBrA4_0101980; **4** (Tc4) = fusion of TcBrA4_0028480 and TcBrA4_0088260; **5** (Tc5) = fusion of TcYC6_0100010 and TcBrA4_0074300; **6** (Tc6) = fusion of TcYC6_0043560 and TcYC6_0122760; **7** (Tc7) = fusion of TcYC6_0083710 and TcBrA4_0130080; **8** (Tc8) = TcYC6_0037170; **9** (Tc11) = TcYC6_0124160; **10** (Tc17) = fusion of TcBrA4_0028230 and TcBrA4_0029760; **11** (Tc19) = TcBrA4_0122270 and TcBrA4_0131050; **12** (tc2-tol2) = TcBrA4_0101960; **13** (3tolt) = portions of TcBrA4_0101970, TcYC6_0077100 and TcYC6_0078140; **14** (beta-tubulin) = TcYC6_0010960; **15** (G10) = TcCLB.504199.20; **16** (Kn107) = TcCLB.508355.250; **17** (LE2) = TcCLB.507447.19; Parvo = Recombinant Canine Parvovirus VP2 (MyBioSource.com).

### Cardiac Troponin

Cardiac troponin (cTnI) analysis was performed using the ADVIA Centaur CP® immunoassay (Ultra-TnI, Siemens Medical Solutions USA, Inc., Malvern, PA) validated in dogs and with a reported range of 0.006 to 50.0 ng/mL [25].

### Statistical analysis

The non-parametric Mann-Whitney U and unpaired t tests of the GraphPad Prism version 9.4 software were used for the analysis of the mouse experimental data. Values are expressed as means ± standard error of mean.

## Ethics Statement

The studies in mice were carried out in strict accordance with the Public Health Service Policy on Humane Care and Use of Laboratory Animals and Association for Assessment and Accreditation of Laboratory Animal Care accreditation guidelines. The protocol was approved by the University of Georgia Institutional Animal Care and Use Committee (AUP #A2021 04-011-Y1-A0). Informed consent was obtained from dog owners prior to their participation, and this study was approved by the Texas A&M University Institutional Committee on Animal Use and Care and the Clinical Research Review Committee (IACUC 2018-0460 CA).

## Results

We had previously shown that weekly [15] or twice weekly (Bustamante, et al, in preparation) administration of high dose (2.5 – 5 X the normal daily dose) BNZ could cure established infections with *T. cruzi* in mice. To determine if BNZ might also prevent the initial establishment of *T. cruzi* in naïve mice, we conducted a pilot study (Figure 1A) using twice weekly delivery of low dose BNZ (100 mg/kg oral, the standard daily dosage for continuous treatment in mice [26]. At 3.5 weeks (between the 5^th^ and 6^th^ BNZ dose) all mice were injected i.p. with 1000 typomastigotes of the luciferase transgenic Colombiana strain of *T. cruzi*. Groups of both wild-type (C57BL/6J) and IFN*γ* deficient mice were included in the study; the IFN*γ* deficient (KO) mice are highly susceptible to *T. cruzi* infection [15, 27] and were expected to reveal even very low levels of persistent infection. Untreated mice in both groups exhibited evidence of active infection at 14 days post-exposure and beyond and as expected, all mice in the IFN*γ* deficient group not receiving prophylaxis experienced uncontrolled infections and had to be euthanized by ∼21 days post-exposure. In contrast, some BNZ-prophylaxed IFN-g KO mice showed low level luciferase signal at day 14 post-exposure but not at 12 weeks, and none of the mice in the BNZ-treated WT group had a detectable luciferase signal above background throughout the study (Fig 1B, C and Fig. S1).

Infection exposure was also confirmed in mice by monitoring the generation of *T. cruzi* -induced CD8^+^ T cells specific for the immunodominant TSKb20 epitope [26]. With the exception of one prophylaxed IFN*γ* KO mouse, all mice exhibited *T. cruzi* -specific CD8^+^ T cell responses with comparable numbers in the untreated WT and prophylaxed IFN*γ* KO mice and lower levels in the prophylaxed WT mice. We have previously shown that the expression of the T cell central memory marker CD127 on these parasite-specific T cells is a useful surrogate for parasite load [15, 26] and the 2 groups under prophylaxis had a substantially higher fraction of CD127^+^ cells in the TSKb20-specific population relative to mice not receiving prophylaxis, consistent with the expected low/absent parasite burden through the 21^st^ week of prophylaxis (Fig 1D).

Twice weekly BNZ was continued for a total of 24 weeks, simulating a seasonal period of vector activity typical of south Texas [13]. In WT mice with or without prophylaxis, infection was not readily detectable by whole animal imaging through the prophylaxis period. However, in the IFN*γ* KO mice, although the infection was largely controlled by the prophylaxis treatment, the systemic infection was barely detectable during the prophylactic treatment period (Fig S1) and within 4 weeks of termination of BNZ prophylaxis, 4 of 5 mice in the previously treated IFN*γ* KO mice group were systemically parasite-positive and had to be euthanized (Figure 1C and Sup Figure 1). The remaining animals in all groups were euthanized at week 33 of the study and skeletal muscle and cardiac tissues were examined for the presence of *T. cruzi* using qPCR. As shown in Fig 1E, all of the untreated WT mice were PCR positive for *T. cruzi* DNA in one or more of the samples from skeletal muscle, heart, or gut. In contrast, at least 3 of the 5 WT mice receiving BZN prophylaxis and the surviving IFN*γ* KO mouse were negative for *T. cruzi* DNA (the remaining 2 WT-treated mice were at or below the level of detection in this assay in skeletal muscle only). Collectively, these results suggest that the prophylaxis regimen employed here in WT mice prevented or truncated an infection in most cases. However, the protective effect of prophylaxis generally required a coincident intact immune response, as most IFN*γ* deficient mice became and remained infected despite prophylaxis.

Building upon these encouraging results in mice, we next asked if BNZ prophylaxis could reduce infection in kennel and hunting dog populations in the south-central U.S. where infection pressure was high, despite other vector control efforts [11]. We conducted an early spring (March to April) screening to identified uninfected dogs in this setting with the intent of capturing infection status prior to the window of high vector activity [13], using a combination of negative serological tests and negative blood PCR. The rationale for using this combination of tests is that although blood PCR can provide solid evidence of active infection, a negative blood PCR assay is not a dependable indicator of absence of infection. Additionally, serologic tests may miss very recent infections. A total of 126 dogs of previously undetermined infection status were screened using a Luminex-based assay previously employed for detection and monitoring *T. cruzi* infection in humans [28-30] and other species [24, 31], a commercial Stat-Pak test (ChemBio, NY) that is validated for humans but commonly used for research purposes in dogs [32, 33], and by blood PCR. Figure 2A displays dogs considered to be infection-negative based upon the Luminex assay and confirmed in most cases by Stat-Pak and indirect fluorescent antibody (IFA) (Texas A&M Veterinary Medical Diagnostic Laboratory, College Station, TX; [32]) assays (Supplemental Table 1; n=57; 45.2% of total screened). Nearly all dogs had detectable antibodies to the parvovirus vaccine antigen, confirming the quality of the serum sample, and in all cases the negative *T. cruzi* serology was supported by a negative blood PCR assay.

**Figure 2.**
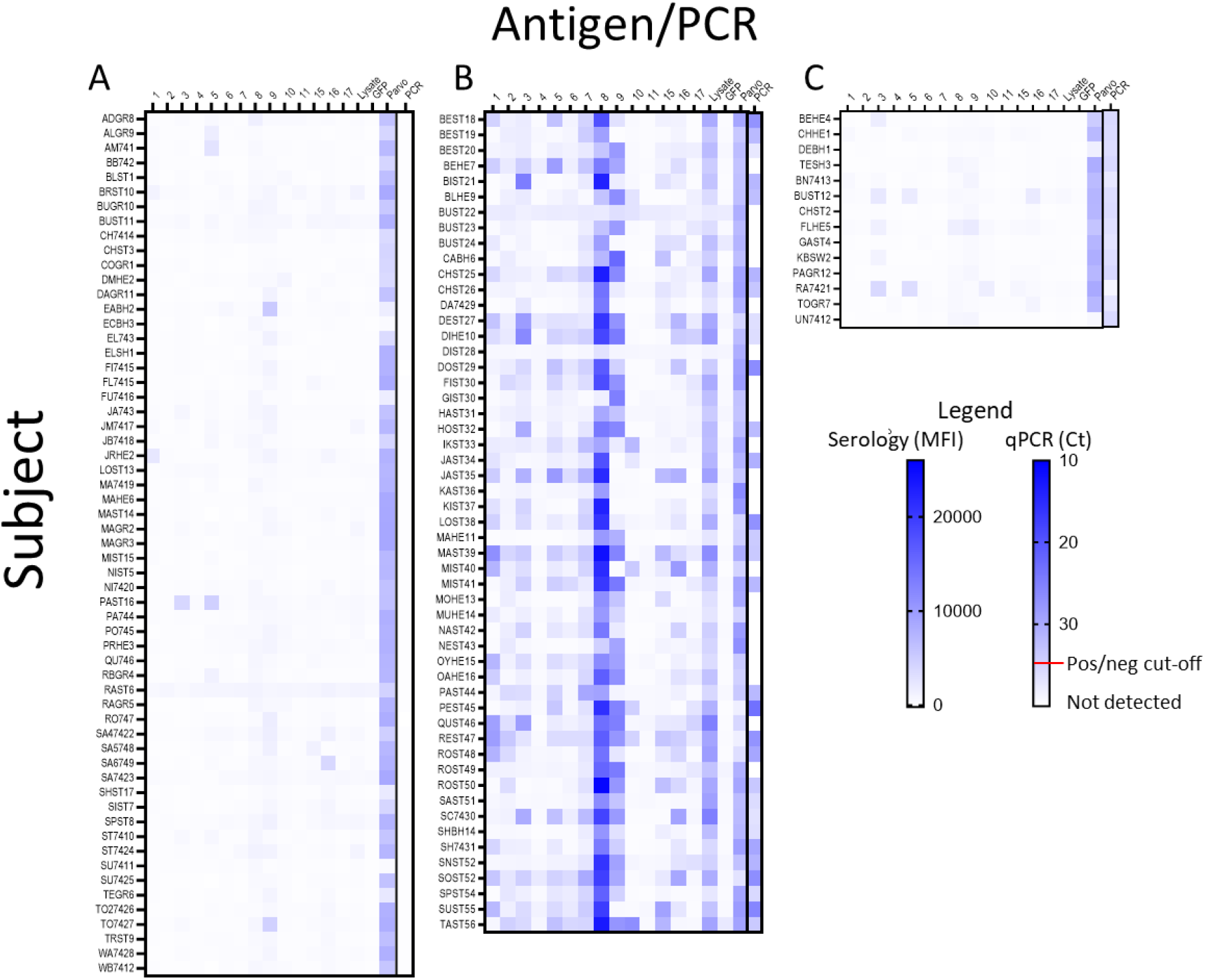
Initial screening results with multiplex serology and PCR to identify *T. cruzi*-negative dogs to enroll in the prophylaxis study. Heatmaps of antibody levels (mean fluorescence intensity; MFI) and qPCR measurement of *T. cruzi* DNA in blood were used to classify dogs as uninfected **(A)**, *T. cruzi*-infected **(B)**, or seronegative but with a blood PCR result that did not reach the cut-off for being considered positive **(C)**. Recombinant *T. cruzi* proteins are defined in the Materials and Methods; lysate = a sonicate of *T. cruzi* trypomastigotes and amastigotes; GFP=recombinant green fluorescent protein (negative antigen control); Parvo = Parvovirus vaccine antigen (positive serological control).

Figure 2B shows the antibody and/or PCR+ dogs that were excluded from the prophylaxis trial by virtue of having prior exposure and likely active infection (n=53; 42.1% of those screened). Roughly half (29/53; 54.7%) of the dogs with detectable (generally robust) serological responses also had *T. cruzi* DNA in their blood as determined by qPCR. It is notable that, as in other species [24, 28-31], the pattern of antigen specificity of antibody responses in these subjects varied extensively, although the antigen in lane 8, a polyubiquitin protein, appeared to be the most reliable. A fraction of the screened dogs (n = 14; 11.1%) were seronegative by Luminex and other assays but had a weak signal in the blood PCR assay that was measurable, but above our standard cutoff for determining infection (Ct < 35.0; Fig 2C and Table S1).

These dogs were initially enrolled in the prophylaxis study as uninfected but 4 of these likely were already infected and these were excluded from prophylaxis study analysis (detailed further below). The 67 animals considered infection-negative and available for enrollment were randomly assigned to either a control (no therapy) or a prophylactic group, which received twice weekly doses of BNZ at a level of not less than 10 mg/kg. This dosage is approximately the normal **daily** dose previously used (mostly ineffectively) to attempt drug-cure in dogs [34-36]. Dogs in these groups were then re-sampled 10-12 weeks (considered the midpoint of the transmission season) and ∼24 weeks (end of season).

Apparently new infections occurred in both the control (n=6; 18.2%) and prophylaxis (n=9; 26.5%) groups, yielding a combined seasonal infection incidence of 22.4%. In all but one case, these infections occurred early in the season, prior to the 12 week screening timepoint. One newly-infected dog in each group was lost to follow up before the 24 week sampling point (one was sold and one died due to *T. cruzi* with severe, acute, lymphohistiocytic, necrotizing pancarditis with intralesional amastigotes). Additionally, one infected dog in each group was removed from the study after the mid-point sampling due to high serum cardiac troponin I (cTnI) levels (indicative of active heart damage) and started on a treatment protocol consisting of higher dose BNZ (to be reported on elsewhere). Thus, BNZ prophylaxis as employed in this study had no impact on the number of new infections in this high intensity transmission setting.

The serial sampling during a transmission season and the stable pattern of the antibody profiles in infected subjects provided the opportunity for several additional novel observations. First, 4 of the 16 seronegative dogs with blood PCR signals exceeding the positive cutoff point (Figure 2C) developed a seropositive profile during follow-up (Figure 4) and 3 of the 4 developed strong blood PCR signals. While these could represent new infections acquired during the follow-up period, they may also indicate dogs that were already infected at the time of the March/April survey but were so early in the infection course that antibody responses had not developed and parasites in the blood were low. The remaining 12 dogs in this subset were equally dispersed in the untreated and prophylaxis groups and none developed evidence of infection during the 24-week follow-up (Figure 3 and Table S1). Looking retrospectively, one dog in each group in the prophylaxis trial was PCR negative (JRHE2, prophylaxis group and PAST16, control group) but was nonetheless also likely already infected at the pre-season survey point, as both had low reactivity to several*T. cruzi* antigens in the multiplex assay at this point but with a pattern that matched the more robust antigen recognition pattern in the Luminex assay that ultimately developed by the 12 and 24 week sampling points (Figure 3).

**Figure 3.**
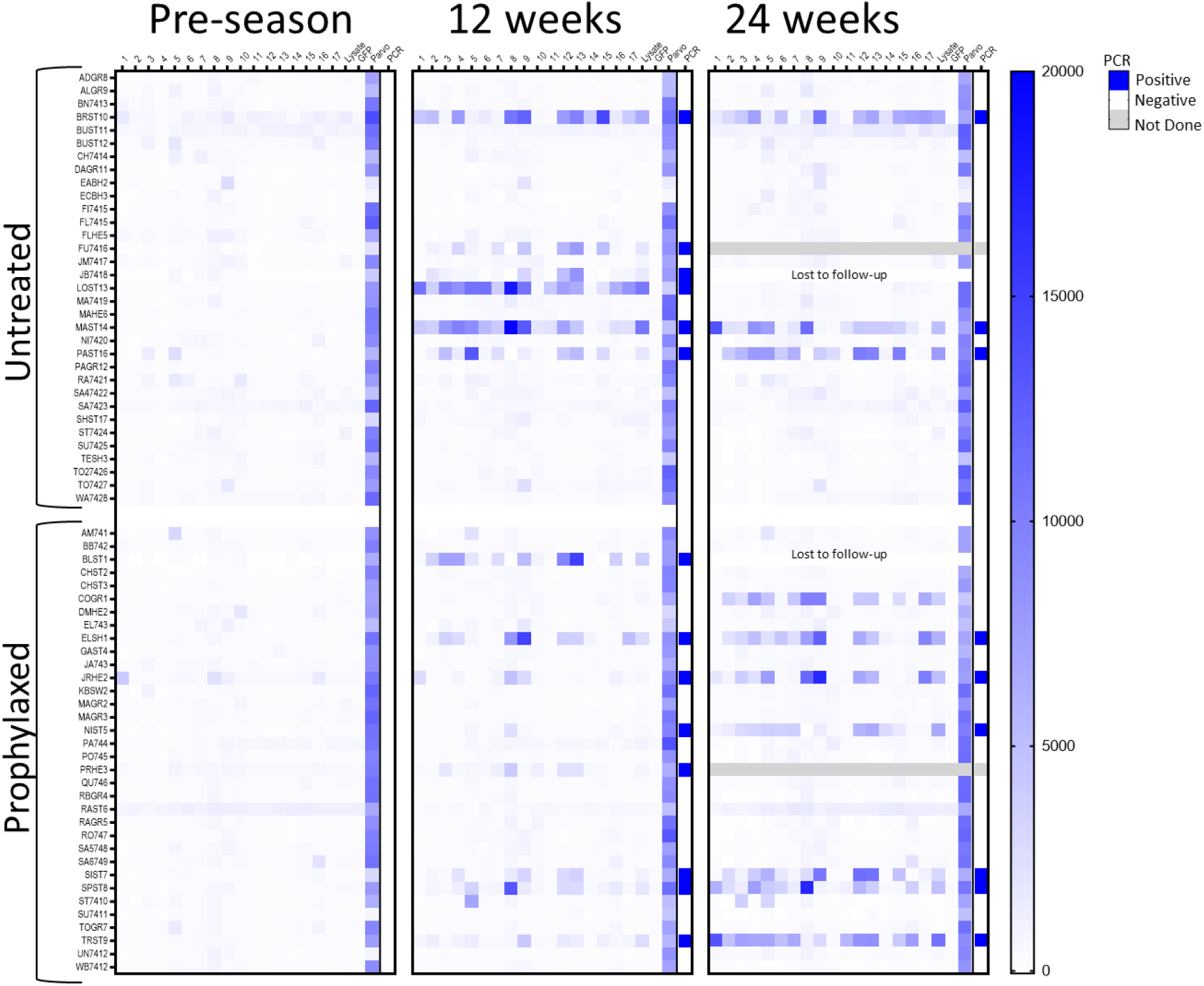
Multiplex serological and blood PCR changes over the transmission season in dogs receiving or not BNZ prophylaxis. Serological and PCR negative dogs (see Figure 2) were randomly assigned to the untreated or BNZ treatment groups and then rescreened at ∼12 and ∼24 weeks after the beginning of prophylaxis. One dog in each group (gray fill) was moved to BNZ treatment protocol after the 12 week sampling point due to high serum cardiac troponin I (cTnI) levels. One dog in each group was lost to follow-up before the 24 week sampling time due to death or sold by owner. Antigens used in the multiplex assay are as described in Figure 2 and the Materials and Methods. Blood PCR was considered positive if below the cut-off Ct value of 35.

**Figure 4.**
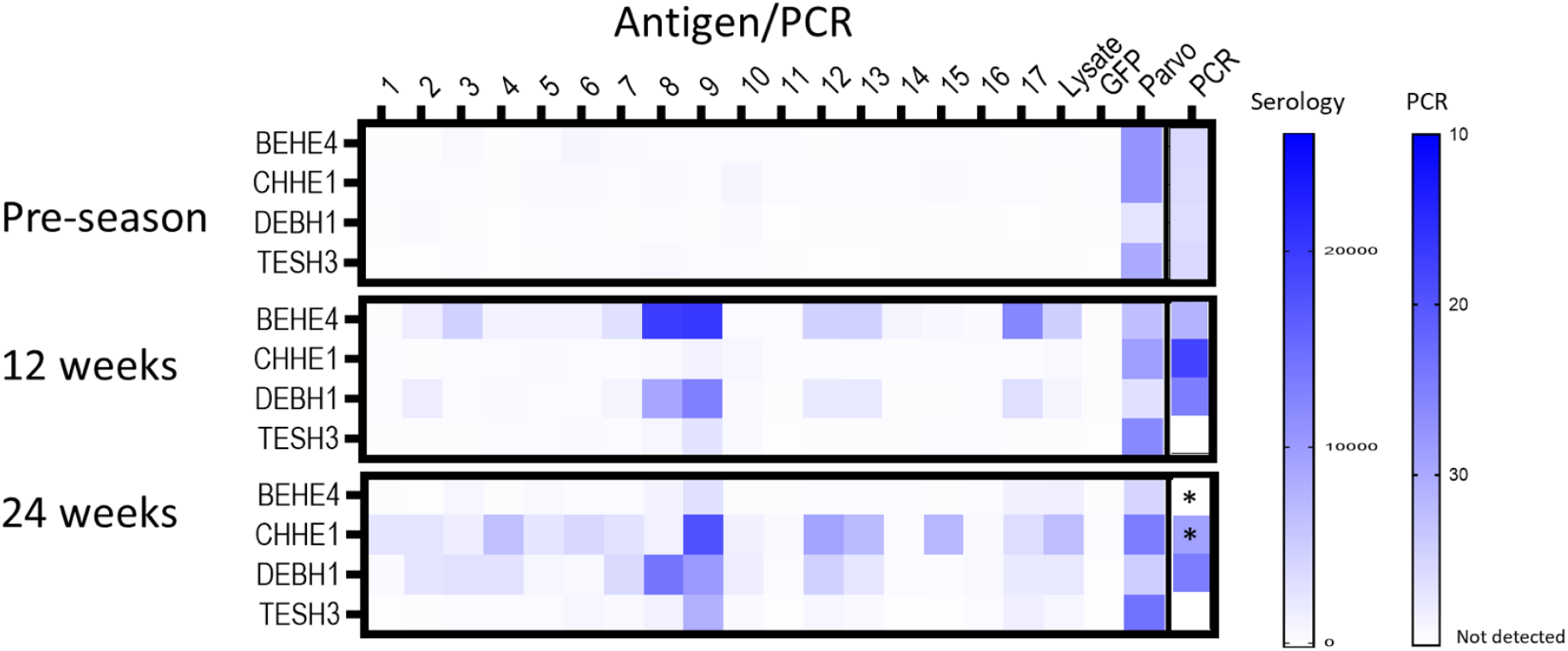
Multiplex serological and blood PCR changes over the transmission season in dogs that were seronegative but PCR positive in the pre-transmission season screening (see Figure 2). Antigens used in the multiplex assay are as described in Figure 2 and the Materials and Methods. * moved to a high dose BNZ treatment protocol after 12 week sample point.

Lastly, one dog in the control group (LOST13, Figure 3) had developed a substantial antibody profile by the 12 week sampling point (at which he was also PCR positive), but had essentially totally lost this response by the 24 week screening point. This dog was also strongly positive by a StatPak antibody test and had detectable serum cTnI (0.021 ng/mL) at 12 weeks but was seronegative and had cTnI below lower detection limit of 0.006 ng/mL at the 24 week sample time point. The continued detection of parvovirus-specific antibodies over this same time course ruled out concerns on sample quality or immune competency. A similar pattern of cure was evident in dog BEHE4 (Figure 4) after he was switched to a BNZ treatment regimen (increasing the dosage to 30 mg/kg twice a week) due to high cTnI levels in the 12 week sample. This dog became PCR negative and seronegative by the 24 week sample point. Collectively, these data support a potential early infection spontaneous cure one dog and BNZ-induced cure in another, both resulting in a rapid decline in serological evidence of the previous infection.

## Discussion

Our previous studies have extensively documented the high risk of *T. cruzi* infection in dogs in south central Texas, including among kenneled working dog populations [11, 21, 32]. New infection rates of up to 30%/year put these valuable animals at high risk for morbidity and early death. It is noteworthy that this high incidence of infection is occurring despite vector awareness education for dog owners and the implementation of a variety of vector control measures. Likely, the abundance of insect vectors and wildlife reservoirs with active *T. cruzi* infections, the outdoor group housing conditions that expose dogs to this infection risk, and the propensity of dogs to ingest these vectors and their feces, all combine to promote this high transmission situation. The lack of interventions such as vaccines that could reduce the incidence of new infections and the high failure rates of potential therapies leave few options for reducing the impact of *T. cruzi* in these settings.

In this study, we explored the potential of BNZ, a drug that has been in use for >50 years for the treatment of *T. cruzi* infection, in a prophylactic modality to attempt to reduce the rate of new infections in dogs. To our knowledge, prophylactic use of trypanocidal drugs has not been previously explored for prevention of new infections with *T. cruzi* [37], although prophylaxis has occasionally been employed to prevent potential exacerbation of infection due to immunosuppression following tissue transplantation in humans [38]. The premise behind these studies was that dosing BNZ at a standard daily dose level, but administering it twice per week instead of daily, could prevent establishment of new infections, or might rapidly terminate those infections before they could become firmly established. The twice-weekly dosing pattern is supported by the ability of BNZ to cure established *T. cruzi* infections in mice using this schedule, albeit with 2.5-5-fold the dose level used here for prophylaxis. Selection of this lower prophylactic dose, bi-weekly schedule was also driven by the practicality in terms of time/effort, cost, and cumulative drug toxicity over a potential 6-month transmission season.

Trials of BNZ-based prophylaxis in laboratory mice yielded promising results, demonstrating a substantial reduction in parasite burden and the ultimate resolution of infection in the majority of immunocompetent mice. Notably, this protocol had much less impact in mice compromised in the ability to produce IFN*γ*, an essential component of the anti-*T. cruzi* immune response. This latter result supports the hypothesis that a competent immune response likely works in concert with trypanocidal drugs to ultimately clear *T. cruzi* infection when treatment is successful.

However, in dogs in a high transmission setting, prophylaxis as applied here failed to prevent new infections during a 6-month transmission season. A number of factors could have contributed to this differential result in mice and dogs, primary among them, the likely differential parasite exposure. Mice received a single sub-lethal intraperitoneal injection of 1000 trypomastigotes, while dogs were potentially exposed repeatedly to infection, possibly at much higher doses, and likely via an oral route. The fact that one dog newly infected during the died during the acute infection and others exhibited high serum cTnI levels, prompting their transfer to a higher treatment dose regimen, supports the higher (and also variable) exposure conditions for dogs in this study. Effective prophylaxis under these scenarios might require a more aggressive dosing regimen, either more frequent or with higher dosage or both.

Although not designed to explicitly address this point, the infection pattern observed in this study supports a seasonality in infection potential. We estimated the transmission season based upon cumulative vector sighting observations collected by our Kissing Bug Community Science initiative [13], which suggested vector activity beginning in May and peaking in July. However, at least 4 dogs were borderline PCR positive but seronegative in early May, and all developed serological responses, 3 of the 4 within 10-12 weeks of the initial sampling. Additionally, two dogs that were PCR-negative, but had a suggestion of a serological response at initial screening, were PCR positive 10-12 weeks later.

Collectively, these data indicate that new infections may be occurring before our early May prophylactic dosing. In combination with the fact that most new infections occurred in the first 12 weeks dosing period, it seems likely that beginning prophylaxis earlier in the year could have had a greater impact in the rate of new infections. Including these four “pre-prophylaxis” infected dogs as among those infected in 2021 year increases the yearly incidence in this setting to 26.8% (19/71).

To our knowledge, no previous study has sampled a susceptible, at risk, animal population at the frequency and with both PCR and multiple serology as employed in this study. In addition to the high infection pressure, this setting is also optimal for collection of novel and valuable data because new dogs are introduced each season, through either breeding or new acquisitions. Among the interesting observations from this first screening year is that 14 of the 15 (or 18 of 19 if the 4 pre-May infections are included) new infections occurred before the calculated mid-point of the estimated transmission season. The reason for this early season bias is not known, but could be due to opportunity based on location, or to behavior (e.g. aggressive bug eaters are more likely to be exposed and become infected) or a combination of the two. Given the high early season incidence, it is clear that the opportunities for infection are high even when bug activity appears low. Identifying the specific links between bug numbers, locations and infection status with the seasonality of new infections in dogs should be addressed in future seasonal studies. Active entomological surveillance at sentinel locations within the geographic region of study would be useful to establish a more precise start of the seasonal insect activity period, so that prophylaxis in future years could be initiated prior to insect activity.

The frequency of sampling applied herein also allowed for the detection of one apparent case of spontaneous cure of an acute infection. Although such spontaneous cures have been anecdotally reported in chronically infected hosts, both animal [39] and human [40], to our knowledge cure during the acute infection has not been documented. The frequency of such cures is worth exploring further and as well, the protection from re-infection that might be afforded by such cases. The very rapid waning of the antibody response in this case, as well as following an apparent BNZ treatment dose-induced cure, was also surprising and would have gone undetected if screening was conducted only once or twice per year, for example.

The use of both of a multiplex serological test and blood PCR to track new infections was critical to the technical success of this project and for revealing some of the less expected observations. The frequent failure of PCR to detect many chronic infections with *T. cruzi* is well recognized [31]. The multiple antigen array used in this study, incorporating >25 parasite proteins, a crude lysate, and both positive and negative control proteins, provides a confidence in detection of infection that is lacking in single antigen or whole parasite assays. However, all serological assays fail infection detection very early, before anti-parasite antibodies have been formed. Thus, the combined use of PCR and multiplex serology provided insights that would have been missed using either approach alone.

Also, not addressed in this study is whether prophylaxis, while not reducing the incidence of infection, might be beneficial by ultimately reducing the severity of disease. Anecdotally we observed that several dogs infected while under prophylaxis nonetheless experienced high serum levels of cTnI, indicating that prophylaxis in these cases was not preventing acute phase disease. A longer-term follow-up of disease development with and without early prophylaxis might be revealing. However, ideally, when infection is detected in these working dogs, they should be enrolled in an effective treatment regimen (as done for dogs BEHE4 and CHHE1) that enhances the chances of resolving the infection and thus preventing disease progression.

Although the seasonal monitoring for new infections in a population at high risk of infection provided new insights into *T. cruzi* transmission and development of immune responses, the prophylaxis approach applied here did not prevent infection in this setting. Future studies might apply prophylaxis earlier in the transmission season or at a more aggressive dosing level. Also, it would be of interest to determine if prophylaxis, while failing to reduce infection incidence, might be beneficial with respect to reducing infection impact either in the short-term, or long-term.

## Acknowledgements

We thank the kennel owners and dog managers, Dr. Madeline Droog, pharmacist, in the Texas A&M University Small Animal Veterinary Medical Teaching Hospital, and Drs. Erin Edwards and Andres de la Concha at Texas A&M Veterinary Medical Diagnostic Laboratory for assistance with the study and Insud/VetPharma, the Mundo Sano Foundation and Humanigen for donation of benznidazole. Facilities in the CTEGD Cytometry Shared Resource Lab were critical to the study. Funding: National Institutes of Health grants R01 AI151148 and R01 AI125738 to RLT; American Veterinary Medical Foundation/Veterinary Pharmacology Research Foundation and University of Texas Southwestern/Texas A&M University Pilot Award, NIH Clinical Translational Science Award (CTSA) 1UL1TR003163 to ABS and SAH; Harry L. Willett Foundation; George A. Robinson Foundation. The funders had no role in study design, data collection and analysis, decision to publish, or preparation of the manuscript.

## Supporting Information Captions

**Supplemental Figure 1**. Images of *T. cruzi* bioluminescent signal in mice (ventral view) at various times post study initiation (see Figure 1A) Note: only 3 of the 5 mice in WT groups were imaged at 24 weeks.

**Supplemental Table 1**. Pre-study dog survey data

