## Supplementary figures and images for "Prophylactic low-dose, bi-weekly benznidazole treatment fails to prevent *Trypanosoma cruzi* infection in dogs under intense transmission pressure"

### Supplemental Figure 1

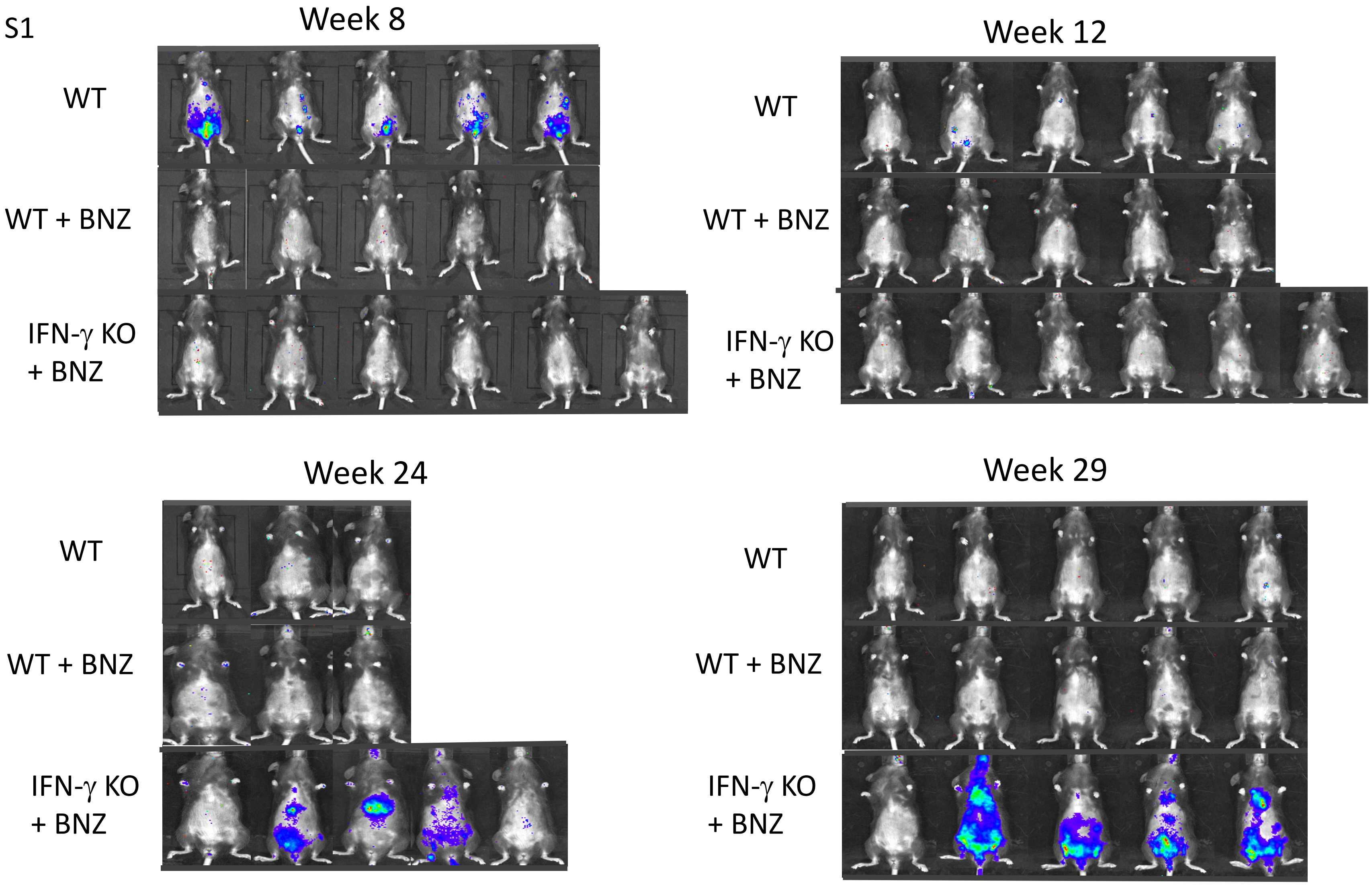
