## Supplemental Table 1 for "Prophylactic low-dose, bi-weekly benznidazole treatment fails to prevent *Trypanosoma cruzi* infection in dogs under intense transmission pressure"

### Pre-study survey data

| Dog ID | Age (years) | Breed | Sex | Assigned group | Serological test results: Luminex | StatPak | Chagas IFA endpoiint titer | Ct value on *T. cruzi* *q* PCR |
|---|---|---|---|---|---|---|---|---|
| **Dogs enrolled as *T. cruzi* -negative** | | | | | | | | |
| ADGR8 | 10.4 | English pointer | female | untreated | Negative | 0 | <20 | nd |
| ALGR9 | 4.3 | English setter | female | untreated | Negative | very faint | <20 | nd |
| AM741 | 2 | English pointer | female | prophylaxed | Negative | 1 | <20 | nd |
| BN7413 | 10 | English pointer | male | untreated | Negative | 0 | <20 | 37.24 |
| BB742 | 6 | English pointer | female | prophylaxed | Negative | 0 | <20 | nd |
| BLST1 | 2 | English pointer | male | prophylaxed | Negative | 0 | <20 | nd |
| BRST10 | 1 | English ponter mix | male | untreated | Negative | 0 | <20 | nd |
| BUST11 | 3 | English pointer | male | untreated | Negative | 0 | <20 | nd |
| BUST12 | 5 | English pointer | male | untreated | Negative | 0 | <20 | 36.06 |
| CH7414 | 6 | English pointer | female | untreated | Negative | very faint | <20 | nd |
| CHST2 | 5 | Labrador retriever | male | prophylaxed | Negative | 0 | <20 | 35.87 |
| CHST3 | 0.25 | English pointer | male | prophylaxed | Negative | 0 | <20 | nd |
| COGR1 | 4.3 | English pointer | male | prophylaxed | Negative | 0 | <20 | nd |
| DMHE2 | 5.75 | Pit bull mix | female spayed | prophylaxed | Negative | 0 | <20 | nd |
| DAGR11 | 4.3 | English pointer | female | untreated | Negative | 0 | <20 | nd |
| EABH2 | unk | Labrador retriever | male | untreated | Negative | 0 | <20 | nd |
| ECBH3 | unk | Labrador retriever | female | untreated | Negative | 0 | <20 | nd |
| EL743 | 10 | English pointer | female | prophylaxed | Negative | 0 | <20 | nd |
| ELSH1 | 0.58 | Border collie | female | prophylaxed | Negative | 0 | <20 | nd |
| FI7415 | 7 | English pointer | male | untreated | Negative | 0 | <20 | nd |
| FL7415 | 2 | English pointer | male | untreated | Negative | 0 | <20 | nd |
| FLHE5 | 5.58 | Black Mouth Cur | female spayed | untreated | Negative | 0 | <20 | 36.46 |
| FU7416 | 0.2 | Cocker Spaniel | male | untreated | Negative | 0 | <20 | nd |
| GAST4 | 0.25 | English pointer | male | prophylaxed | Negative | 0 | <20 | 36.93 |
| JA743 | 11 | English pointer | male | prophylaxed | Negative | very faint | <20 | nd |
| JM7417 | 11 | English pointer | male | untreated | Negative | 0 | <20 | nd |
| JB7418 | 0.2 | Cocker Spaniel | male | untreated | Negative | 0 | <20 | nd |
| JRHE2 | unk | Black Mouth Cur mix | male, neutered | prophylaxed | Negative | 0 | <20 | nd |
| KBSW2 | 1.5 | Border collie | male | prophylaxed | Negative | 0 | <20 | 35.94 |

| | | | | | | | | |
|---|---|---|---|---|---|---|---|---|
| LOST13 | 2 | English pointer | female | untreated | Negative | 0 | <20 | nd |
| MA7419 | 5 | English pointer | female | untreated | Negative | very faint | <20 | nd |
| MAHE6 | 1.25 | Great Pyrenees | female spayed | untreated | Negative | 0 | <20 | nd |
| MAST14 | 2 | German Shorthair Pointer | female | untreated | Negative | 0 | <20 | nd |
| MAGR2 | 4 | German Shorthair Pointer | male | prophylaxed | Negative | very faint | <20 | nd |
| MAGR3 | 3.3 | English pointer | female | prophylaxed | Negative | 0 | <20 | nd |
| NIST5 | 0.25 | English pointer | female | prophylaxed | Negative | 0 | <20 | nd |
| NI7420 | 11 | Labrador retriever | female | untreated | Negative | 0 | <20 | nd |
| PAST16 | 5 | English pointer | female | untreated | Negative | very faint | <20 | nd |
| PAGR12 | 6.3 | English pointer | male | untreated | Negative | 0 | <20 | 35.37 |
| PA744 | 11 | English pointer | female | prophylaxed | Negative | very faint | <20 | nd |
| PO745 | 4 | Cocker Spaniel | female | prophylaxed | Negative | 0 | <20 | nd |
| PRHE3 | 3.16 | Pit bull terrier | female spayed | prophylaxed | Negative | 0 | <20 | nd |
| QU746 | 5 | English pointer | female | prophylaxed | Negative | 0 | <20 | nd |
| RBGR4 | 2.8 | English Cocker Spaniel | male | prophylaxed | Negative | very faint | <20 | nd |
| RAST6 | 4 | English setter | male, neutered | prophylaxed | Negative | 0 | <20 | nd |
| RA7421 | 2 | English pointer | male | untreated | Negative | 0 | <20 | 38.87 |
| RAGR5 | 1.6 | English Cocker Spaniel | male | prophylaxed | Negative | 0 | <20 | nd |
| RO747 | 1.75 | Labrador retriever | male | prophylaxed | Negative | very faint | <20 | nd |
| SA47422 | 10 | Pointer cross | female | untreated | Negative | very faint | <20 | nd |
| SA5748 | 8 | English pointer | female | prophylaxed | Negative | 0 | <20 | nd |
| SA6749 | 10 | English pointer | female | prophylaxed | Negative | 0 | <20 | nd |
| SA7423 | 5 | English pointer | male | untreated | Negative | 0 | <20 | nd |
| SHST17 | 1 | Labrador retriever | male | untreated | Negative | 0 | <20 | nd |
| SIST7 | 2 | English setter | female | prophylaxed | Negative | 0 | <20 | nd |
| SPST8 | 5 | English pointer | female | prophylaxed | Negative | 0 | <20 | nd |
| ST7410 | 7 | Labrador retriever | female | prophylaxed | Negative | 0 | <20 | nd |
| ST7424 | 6 | German Shorthair Pointer | male | untreated | Negative | 0 | <20 | nd |
| SU7411 | 0.2 | Cocker Spaniel | female | prophylaxed | Negative | 0 | <20 | nd |
| SU7425 | 2 | English pointer | female | untreated | Negative | 0 | <20 | nd |
| TEGR6 | 4.3 | English pointer | male | prophylaxed | Negative | 0 | <20 | nd |
| TO27426 | 3 | Cocker Spaniel | male | untreated | Negative | 0 | <20 | nd |
| TO7427 | 2 | English pointer | male | untreated | Negative | 0 | <20 | nd |
| TOGR7 | 6.1 | English pointer | male | prophylaxed | Negative | 0 | <20 | 36.92 |

| | | | | | | | | |
|---|---|---|---|---|---|---|---|---|
| TRST9 | 2 | Labrador retriever | male | prophylaxed | Negative | 0 | <20 | nd |
| UN7412 | 0.2 | Cocker Spaniel | male | prophylaxed | Negative | very faint | <20 | 35.45 |
| WA7428 | 3 | Cocker Spaniel | male | untreated | Negative | very faint | <20 | nd |
| WB7412 | 2 | English pointer | female | prophylaxed | Negative | 0 | <20 | nd |
| **Dogs potentially infected at time of pre-study screen** | | | | | | | | |
| BEHE4 | 6 | Black Mouth Cur | male | withdrawn | Negative | 0 | <20 | 35.52 |
| CHHE1 | 8 | Labrador retriever | male | withdrawn | Negative | 0 | <20 | 35.65 |
| DEBH1 | 9 | Labrador retriever | female | withdrawn | Negative | 0 | <20 | 36.04 |
| TESH3 | 1.5 | Kelpie | male | withdrawn | Negative | 0 | <20 | 35.45 |
| **Dogs *T. cruzi* -positive** | | | | | | | | |
| BEST18 | 2 | English pointer | female | not enrolled | Positive | 4 | not tested | 27.19 |
| BEST19 | 2 | English pointer | female | not enrolled | Positive | 3 | not tested | 31.8 |
| BEST20 | 4 | English pointer | female | not enrolled | Positive | 3 | not tested | 37.19 |
| BEHE7 | 6.5 | Catahoula mix | female spayed | not enrolled | Positive | 3 | 640 | nd |
| BIST21 | 16 | English pointer | male, neutered | not enrolled | Positive | 4 | not tested | 31.48 |
| BLHE9 | 9.83 | Catahoula | male | not enrolled | Positive | 2 | 1280 | 32.46 |
| BUST22 | 4 | English pointer | male | not enrolled | Positive | 1 | not tested | nd |
| BUST23 | 8 | English setter | male | not enrolled | Positive | 1 | not tested | nd |
| BUST24 | 6 | Labrador retriever | male | not enrolled | Positive | 3 | not tested | nd |
| CABH6 | unk | Labrador retriever | female | not enrolled | Positive | 2 | 320 | nd |
| CHST25 | 8 | English pointer | male | not enrolled | Positive | 4 | not tested | 31.06 |
| CHST26 | 5 | German Shorthair Pointer | male | not enrolled | Positive | 1 | not tested | 33.21 |
| DA7429 | 10 | English pointer | male, neutered | not enrolled | Positive | 4 | 320 | nd |
| DEST27 | 2 | English pointer | female | not enrolled | Positive | 4 | not tested | 36.18 |
| DIHE10 | 8.16 | Pit bull terrier | male | not enrolled | Positive | 3 | 5120 | 35.97 |
| DIST28 | 5 | German Shorthair Pointer | female | not enrolled | Negative | 1 | not tested | nd |
| DOST29 | 8 | English pointer | female | not enrolled | Positive | 1 | not tested | 26.36 |
| FIST30 | 7 | English pointer | female | not enrolled | Positive | 3 | not tested | nd |
| GIST30 | 6 | English pointer | female | not enrolled | Positive | 3 | not tested | nd |
| HAST31 | 9 | English pointer | male | not enrolled | Positive | 3 | not tested | nd |
| HOST32 | 9 | English pointer | male | not enrolled | Positive | 3 | not tested | 31.9 |
| IKST33 | 5 | English pointer | male | not enrolled | Positive | 3 | not tested | nd |

| | | | | | | | | |
|---|---|---|---|---|---|---|---|---|
| JAST34 | 9 | German Shorthair Pointer | female | not enrolled | Positive | 3 | not tested | 30.69 |
| JAST35 | 2 | Labrador retriever | female | not enrolled | Positive | 4 | not tested | nd |
| KAST36 | 5 | English pointer | female | not enrolled | Positive | 2 | not tested | nd |
| KIST37 | 5 | English pointer | female | not enrolled | Positive | 1 | not tested | nd |
| LOST38 | 2 | English pointer | female | not enrolled | Positive | 4 | not tested | 27 |
| MAHE11 | 6 | Pit bull mix | female spayed | not enrolled | Positive | 3 | 640 | 33.77 |
| MAST39 | 11 | English pointer | male | not enrolled | Positive | 2 | not tested | 33.56 |
| MIST40 | 3 | unk | male | not enrolled | Positive | 2 | not tested | nd |
| MIST41 | 5 | English pointer | female spayed | not enrolled | Positive | 4 | not tested | 31.56 |
| MOHE13 | 4 | Mixed breed | female | not enrolled | Positive | 3 | 320 | nd |
| MUHE14 | 6.5 | Catahoula mix | female spayed | not enrolled | Positive | 2 | 1280 | nd |
| NAST42 | 5 | English pointer | male | not enrolled | Positive | 3 | not tested | nd |
| NEST43 | 7 | English pointer | female | not enrolled | Positive | 1 | 2560 | nd |
| OYHE15 | 6 | Pit bull mix | male | not enrolled | Positive | 3 | 2560 | nd |
| OAHE16 | 7 | Hound mix | female spayed | not enrolled | Positive | 3 | 2560 | nd |
| PAST44 | 6 | English setter | female | not enrolled | Positive | 3 | not tested | 31.71 |
| PEST45 | 8 | Labrador retriever | female | not enrolled | Positive | 2 | not tested | 23.2 |
| QUST46 | 10 | English pointer | female | not enrolled | Positive | 3 | not tested | nd |
| REST47 | 7 | English pointer | male | not enrolled | Positive | 3 | not tested | 27.23 |
| ROST48 | 10 | English pointer | male | not enrolled | Positive | 4 | not tested | 29.51 |
| ROST49 | 3 | English pointer | male | not enrolled | Positive | 3 | not tested | nd |
| ROST50 | 8 | English pointer | female | not enrolled | Positive | 3 | not tested | 33.44 |
| SAST51 | 4 | Labrador retriever | female | not enrolled | Positive | 4 | not tested | 35.84 |
| SC7430 | unk | Cocker Spaniel | male | not enrolled | Positive | 3 | 2560 | 36.58 |
| SHBH14 | unk | Labrador retriever | male | not enrolled | Positive | 3 | 320 | 36.14 |
| SH7431 | 2 | English pointer | female | not enrolled | Positive | 1 | 160 | 29.43 |
| SNST52 | 8 | English pointer | female spayed | not enrolled | Positive | 2 | not tested | 31.11 |
| SOST52 | 9 | English pointer | male | not enrolled | Positive | 2 | not tested | 26.66 |
| SPST54 | 8 | German Shorthair Pointer | male | not enrolled | Positive | 4 | not tested | 35.49 |
| SUST55 | 3 | English pointer | female | not enrolled | Positive | 2 | not tested | 25.68 |
| TAST56 | 4 | English pointer | male | not enrolled | Positive | 2 | not tested | 33.76 |

nd = not detected

unk = unknown
